# GITAR: an Open Source Tool for Analysis and Visualization of Hi-C Data

**DOI:** 10.1101/259515

**Authors:** Riccardo Calandrelli, Qiuyang Wu, Jihong Guan, Sheng Zhong

**Author notes:** Equal contribution (Riccardo Calandrelli, Qiuyang Wu). Corresponding author. (Calandrelli R).

## Abstract

Interactions between chromatin segments play a large role in functional genomic assays and developments in genomic interaction detection methods have shown interacting topological domains within the genome. Among these methods, Hi-C plays a key role. Here, we present GITAR (Genome Interaction Tools and Resources), a software to perform a comprehensive Hi-C data analysis, including data preprocessing, normalization, visualization and topologically associated domains (TADs) analysis. GITAR is composed of two main modules: 1) HiCtool, a Python library to process and visualize Hi-C data, including TADs analysis and 2) Processed data library, a large collection of human and mouse datasets processed using HiCtool. HiCtool leads the user step-by-step through a pipeline which goes from the raw Hi-C data to the computation, visualization and optimized storage of intra-chromosomal contact matrices and topological domain coordinates. A large collection of standardized processed data allows to compare different datasets in a consistent way and it saves time of work to obtain data for visualization or additional analyses. GITAR enables users without any programming or bioinformatic expertise to work with Hi-C data and it is freely available for the public at http://genomegitar.org as an open source software.

## Introduction

Genomes are more than linear sequences, with DNA folding-up into elaborate physical structures that allow for extreme spatial compactness of the genetic material and play also an important role in epigenetic regulation [1, 2]. During the past fifteen years, several techniques have been developed to explore the architecture of genomes, such as Chromosome Conformation Capture (3C) [3], Circular Chromosome Conformation Capture (4C) [4], Chromosome Conformation Capture Carbon Copy (5C) [5], Hi-C [6] and ChIA-PET [7]. Among these techniques, Hi-C is one of the most important and it allowed for the first time a genome-wide mapping of chromatin interactions. In 2012, Dixon et al. [8] exploited Hi-C data to identify large, megabase-sized local chromatin interaction domains that they named “topological domains”. These domains appeared widespread along the genome, conserved between mice and humans and stable across different cell types. Later in 2014, Hi-C was improved by performing the protocol in intact nuclei (*in situ* Hi-C), to generate higher resolution contact maps [9].

In order to process Hi-C data, several steps are required. First, raw data are subjected to preprocessing including read pairs alignment and filtering to remove low quality mapped reads, PCR duplicates and non-informative pairs. In addition, some strategies could be implemented to improve the mapping outcome [10]. Next, contact matrices are generated by partitioning the linear genome into loci of a certain size, that correspond to the rows and columns of the contact matrix. Given that, each entry of the matrix contains the number of contacts observed between the two corresponding loci. Contact matrices are then normalized, to remove major systematic biases introduced in the experiment. These matrices are usually visualized using heatmaps, with pixel intensity proportional to the number of contacts observed between the corresponding pair of loci. To accomplish these tasks, we developed HiCtool, a pipeline to process and visualize Hi-C data, including also topologically associated domains (TADs) analysis. Moreover, we successfully processed and made available to the public a large collection of human and mouse datasets of different cell lines and conditions, to allow consistent comparison or custom additional analyses.

Here, we present GITAR (Genome Interaction Tools and Resources), a standardized, easy and flexible solution to manage Hi-C genomic interaction data, from processing to storage and visualization, composed of the two modules mentioned above: HiCtool and the processed data library. Having one comprehensive tool to perform the analysis has the key advantage of avoiding the difficulty related to data integration, when different tasks are performed by different software and the input data are required to be in a specific format for each of them.

## Implementation

### HiCtool: a standardized pipeline to process and visualize Hi-C data

HiCtool is an open-source bioinformatic tool based on Python, which integrates several software to perform a standardized Hi-C data analysis, from the raw data to the visualization of intra-chromosomal heatmaps and the identification of TADs. We implemented a pipeline divided into three main sections: data preprocessing, data analysis and visualization, and topological domain analysis (**Figure 1**). HiCtool documentation and code are available at http://doc.genomegitar.org.

**Figure 1.**
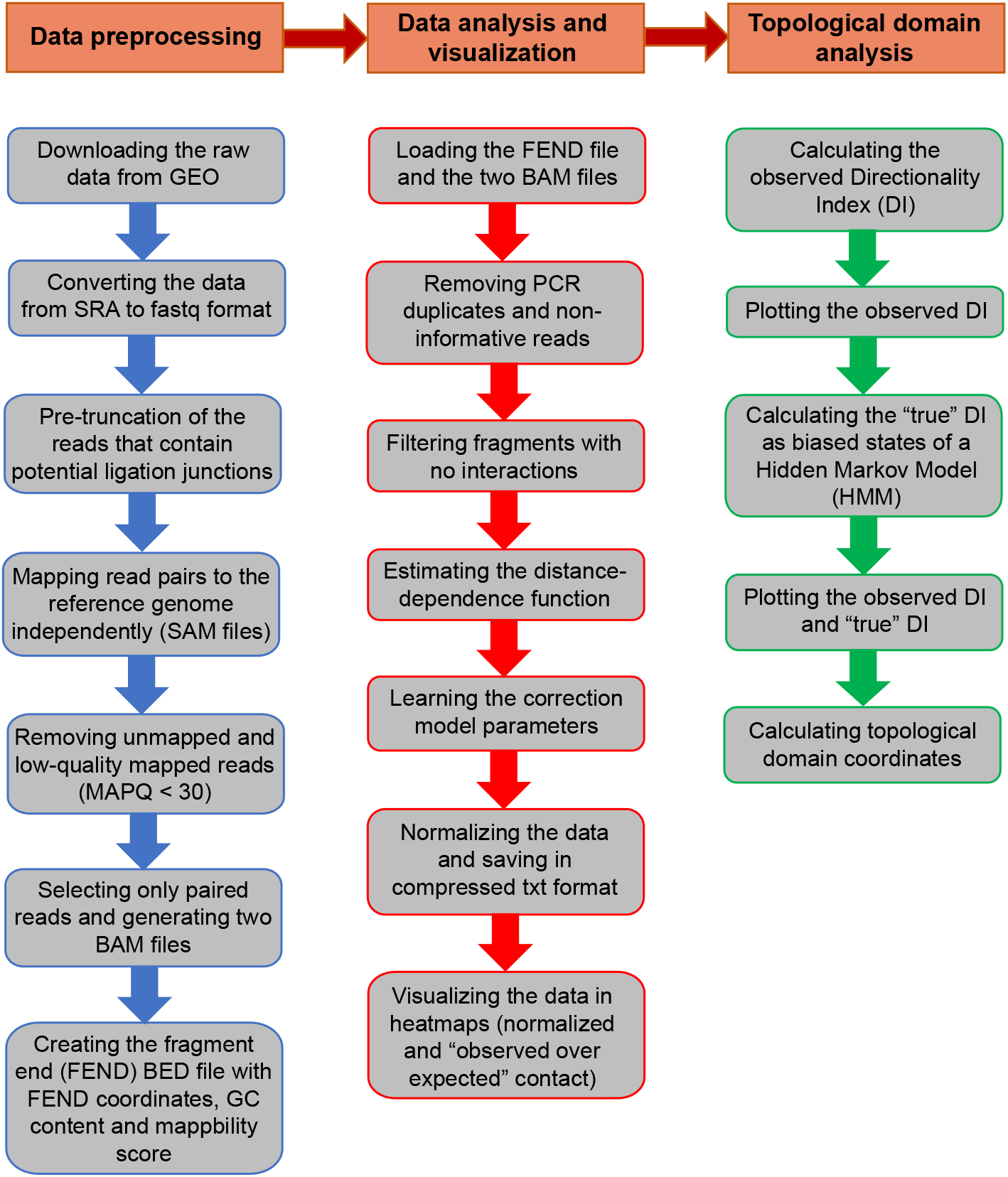
HiCtool workflow schema. HiCtool is a pipeline divided into three main sections: data preprocessing, data analysis and visualization to normalize the contact data and plot heatmaps, and topological domain analysis to calculate Directionality Index (DI), “true” DI using a Hidden Markov Model and topological domain coordinates.

The data preprocessing pipeline takes files downloaded in SRA format and it contains several steps to generate input files for the normalization procedure. To do so, HiCtool integrates Python code, Unix code and several software including SRA Toolkit, Bowtie 2, SAMTools and BEDTools. The preprocessing steps include: downloading the SRA raw data and conversion to fastq format (SRA Toolkit); pre-truncation of the reads that contain potential ligation junctions [10] (Python code); mapping of the read pairs over the reference genome independently to avoid any proximity constraint (Bowtie 2); filtering out unmapped and low-quality mapped reads (MAPQ < 30), then selecting only paired reads (SAMTools and Unix code); creating a fragment end (FEND) file used to normalize the data (Bowtie 2, SAMTools, BEDTools, Python code and Unix code). This file contains restriction sites coordinates and additional information such as GC content and mappability score of the fragment ends. The generation of the FEND file was optimized using parallelized computation to sensibly reduce the time complexity. The outputs of the data preprocessing section are two BAM files related to the first and second read in the pairs and the FEND file in BED format.

The data analysis and visualization pipeline provides the code to normalize the data and plot the contact heatmaps. The complex experimental Hi-C protocol unavoidably produces several technical biases. According to Yaffe and Tanay, 2011 [11], these biases are related to spurious ligation products between fragments, and fragment length, GC content and mappability. Spurious ligation products generate paired-reads whose sum of the two distances to the nearest restriction sites is larger than 500 bp. About fragment length, long and short fragments may have a different ligation efficiency. The probability of contact can be also influenced by the GC content near the ligated fragment ends, up to 200 bp next to the restriction sites, and by the mappability of the sequence, up to 500 bp next to the restriction sites. In addition, it has been observed that transcription start sites (TSSs) and CTCF binding sites influence the frequency of interactions, due to a specific local chromosomal architecture around them [11]. Analysis of the distribution of *cis* contacts involving fragment ends located 0-5 kb upstream of an active TSS showed a strong enrichment 20 kb to ~400 kb upstream and downstream, increasing the probability that long-range contacts may be associated with the active transcriptional state. In addition, these fragment ends produce an asymmetry of the *cis*-interaction profile near the TSS. CTCF binding sites are involved in chromatin organization by providing demarcation for topological domains. This gives rise to a *cis* contact asymmetry of fragments located 0-5 kb on one side of a CTCF-binding site over a range up to ~400 kb, confirming the correlation between CTCF and local chromosomal domains. To normalize the data, we used the Python package HiFive [12]. This allowed to remove all the previous mentioned technical biases taking also into account of “biological biases” that influence the contact frequency (TSSs and CTCF-bound sites), to do not confound technical artifacts with meaningful biological features while learning correction parameters. Before normalization, PCR duplicates and non-informative reads, produced by the possibility of incomplete restriction enzyme digestion and fragment circularization, were removed. Moreover, HiFive allowed to estimate the distance-dependence signal from the data before normalization, in order to avoid biases caused by the restriction site distribution. Restriction sites over the genome are unevenly distributed and this results in many different distances between fragments and their neighbors. Since the interaction frequency is strongly inversely-related to the inter-fragment distance, this means that fragments surrounded by shorter ones will show higher nearby interaction values than those with longer adjacent fragments [12]. To learn the contact correction parameters associated to fragment features, we used the explicit-factor correction scheme of Yaffe and Tanay [11], performed by the HiFive Binning algorithm, which has a consistent performance across all binning resolutions [12]. Given the features of the input FEND file, the adjustment derived by the likelihood optimization procedure can be explicitly attributed to the cross-correlation of fragment length, GC content and mappability. “Biological biases” associated to TSSs and CTCF binding sites are considered at this step by excluding fragments interacting within a distance of 500 kb from the model. After having learned the correction parameters, for any arbitrary division of the genome into bins, we computed two matrices for each chromosome: an observed intra-chromosomal contact matrix *O[i,j]*, where each entry contains the observed read count between the regions identified by the bins *i* and *j*, and a correction matrix *E[i,j]*, where each entry contains the sum of corrections for the read pairs between bins *i* and *j*. Then, the normalized contact matrix *N[i,j]* is calculated using the formula:

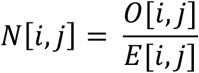

where each entry contains the corrected contact count according to the previous model. We computed also the “observed over expected” contact matrix, where the expected counts are calculated considering both the learned correction parameters and the linear distance between read pairs, given the property that the average intrachromosomal contact probability for pairs of loci decreases monotonically with increasing of their linear genomic distance [6]. Finally, the pipeline allows to plot the heatmaps, with additional histogram of the data distribution (**Figure 2**). The intensity of each pixel in the normalized heatmap is the contact count between the corresponding loci (Figure 2A). For the “observed over expected” (O/E) heatmap, the intensity of each pixel represents the log_2_ of the O/E contact count, to allow easier interpretation of contact enrichment or depletion (Figure 2B). For the visualization functionality, only the Python library Matplotlib was used, without any other external software integration.

**Figure 2.**
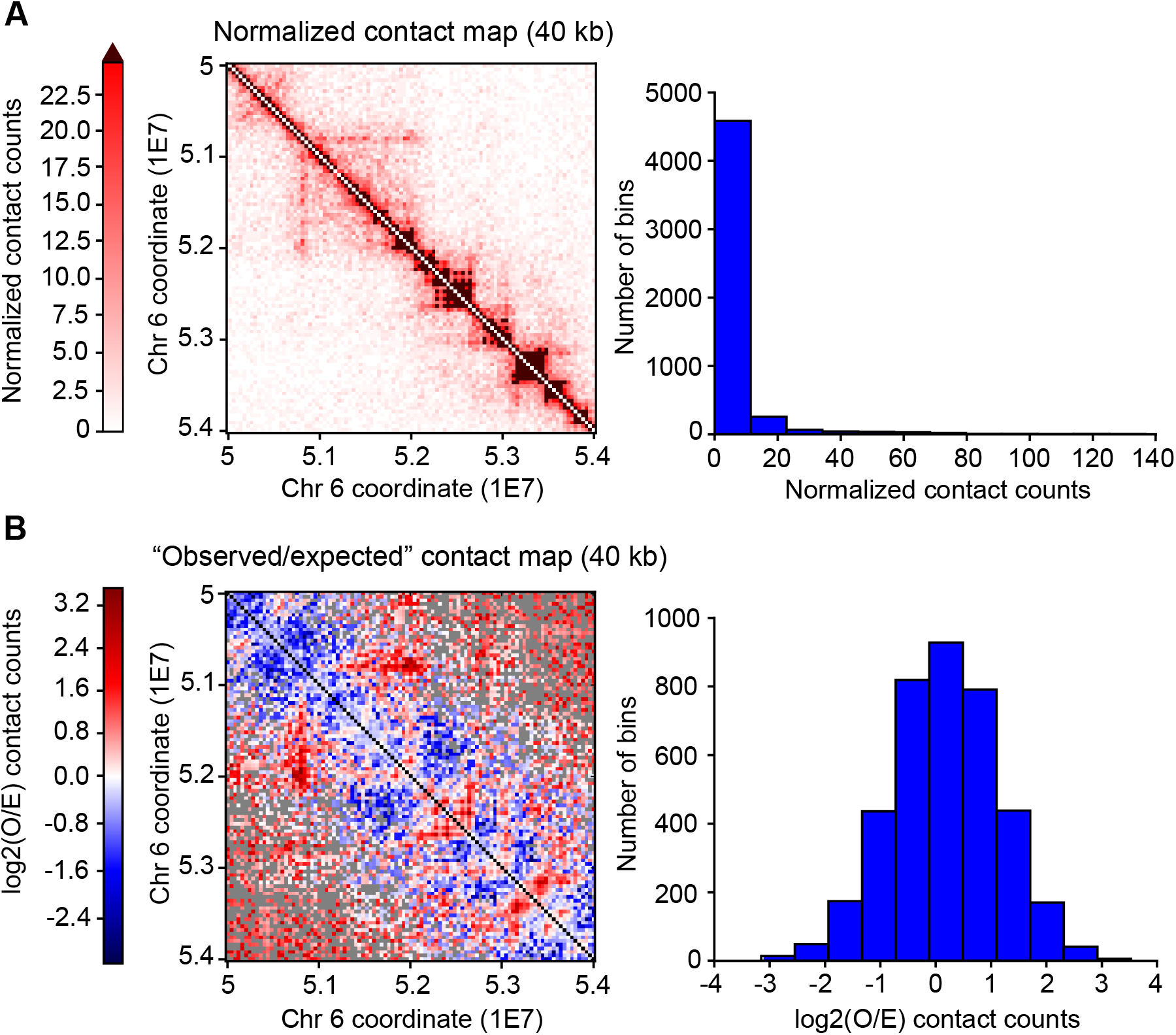
Contact heatmaps for chromosome 6: [50-54] Mb at 40 kb resolution. **A**. Normalized contact matrix, where each entry contains the corrected contact count. The upper limit of the colormap is the 95^th^ percentile of the contact counts; values above this limit are represented in brown in the heatmap. This allows to emphasize local chromatin structures (TADs and sub-TADs) over the background. The histogram displays the normalized contact count distribution, as number of contacts (x-axis) showed in how many bins of the contact matrix (y-axis). Range from 0 to 137 contacts. **B**. “Observed over expected” (O/E) contact matrix, where each entry contains the log_2_ of the observed over expected contact counts, considering the linear distance between loci and the learned correction parameters. Black pixels are for loci with no expected contacts; gray pixels are for loci with no observed contacts. The histogram displays the “observed over expected” (O/E) contact count distribution, as log_2_(O/E) contact counts (x-axis) showed in how many bins of the contact matrix (y-axis). Range from −3.153 to 3.530.

The topological domain analysis section provides the code to calculate the Directionality Index (DI) and the topological domain coordinates [8] from the normalized contact data. In order to do so, the observed DI is used to calculate the “true” DI using a Hidden Markov Model (HMM), implemented with the Python package hmmlearn. Both the observed DI and the “true” DI values can be plotted in the same figure (**Figure 3**), therefore it is possible to directly infer about the presence of topological domains and boundaries over the genomic region under analysis. Topological domain coordinates are then calculated using the shifts of the HMM biased states according to Dixon et al. [8]. A domain is initiated at the beginning of a single downstream biased HMM state and it is continuous throughout any consecutive downstream biased state; the domain will then end when the last in a series of upstream biased states is reached. The entire topological domain analysis pipeline is programmed in Python.

**Figure 3.**
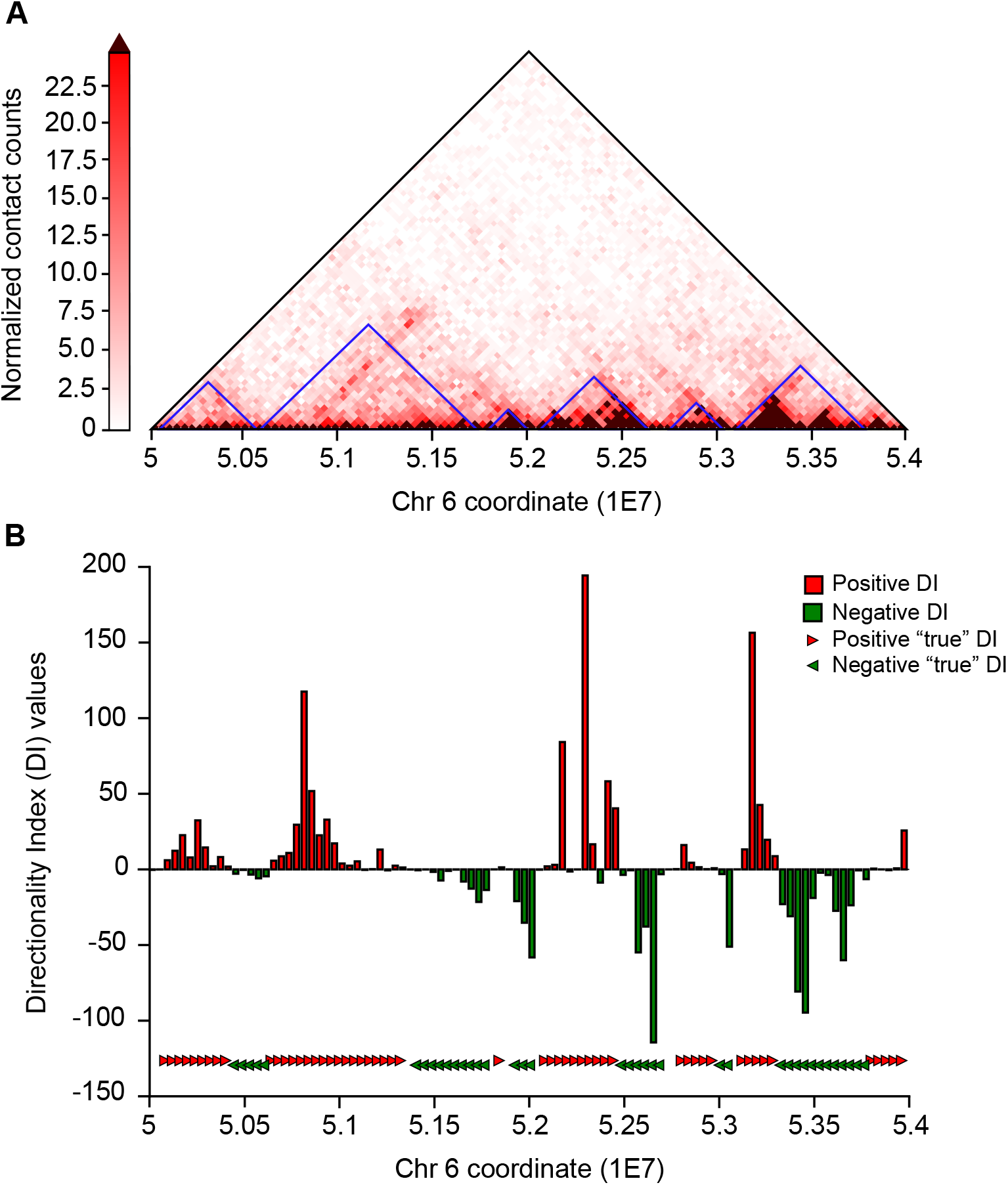
Topological domains for chromosome 6: [50-54] Mb at 40 kb resolution. **A**. Triangular part of the normalized contact matrix. Topological domains calculated with GITAR are plotted in blue along the diagonal. **B**. Directionality Index (DI) values and “true” DI values (HMM biased states). Positive “true” DI and negative “true” DI refer to downstream and upstream HMM biased states respectively. HMM biased states are plotted to show the correspondence with topological domain boundaries (y-axis is not informative in this case). According to the HMM state shifts, six topological domains and seven topological domain boundaries are present.

Here, we showed the capability of GITAR to handle high-resolution Hi-C data by processing and visualizing the *in situ* Hi-C dataset from Rao et al., 2014 [9] (GEO accession number: GSM1551550). For the normalization pipeline, this capability was stated in Sauria et al., 2015 [12], where the authors demonstrated the ability of HiFive to efficiently handle high-resolution data.

### Processed data library

The processed data library is a collection of standardized processed datasets using HiCtool, available for the public at http://data.genomegitar.org. We have already run 19 datasets of human (*hg38*), taken from the library on the 4D Nucleome (4DN) Web Portal (https://4dnucleome.org) [13], and 2 datasets of mouse (*mm10*). The 4DN library is a collection of genome interaction papers related to the Chromosome Conformation Capture based assays (3C, 4C, 5C and Hi-C). Specifically, for GITAR we referred only to Hi-C derived datasets.

Four different outputs for each chromosome of a processed dataset are computed and saved to file: contact matrices, DI values, HMM biased states (“true” DI values) and topological domain coordinates. All the outputs are in txt format, and the functions to load each type of data are provided. Here, we processed the data at 40 kb resolution to perform topological domain analysis according to Dixon et al. [8]. Already at this resolution, contact matrices contain several million of elements per each chromosome, requiring big storage space and relatively high data saving and loading time. To reduce storage usage and computing time, we used a compressed format to save contact matrices. For the normalized contact data from Dixon et al., 2012 [8] (GEO accession number: GSM862723) at the resolution of 40 kb, 1.3 GB of storage was required if the full contact data are saved, only 97 MB using our method. This means that storage usage was reduced of 92% (58% if the files are compressed). About the computing time, saving and loading full contact matrices required respectively 6 minutes and 3 minutes (2.9 GHz Intel Core i5, 16 GB of RAM). Saving and loading our parsed data required 1 minute for both. This means 83% and 67% of data saving and loading time reduction respectively. Conversely, domain coordinates are in a format to be read directly. Each line of the txt file refers to a topological domain, with tab separated start and end coordinates, allowing easy access and readability.

## Authors’ contribution

RC and SZ conceived the project. RC and QW developed the software and performed the analysis. RC wrote the paper, QW and JG contributed to paper writing. All authors read and approved the final manuscript.

## Acknowledgements

This work was supported by the National Institutes of Health of USA (Grant No. U01CA200147 and Grant No. DP1HD087990) awarded to SZ.

## Supplementary text

HiCtool is an open source software and the source code and documentation are available at doc.genomegitar.org. The preprocessing is based on Python code, Unix code and several software (SRA Toolkit, Bowtie 2, SAMTools and BEDTools). The rest of the analysis is programmed in Python, therefore tasks can be performed with single function calls or script executions.

### Mapping

We performed pre-truncation on reads that contain potential ligation junctions before mapping (Ay et al., 2015). All the reads were preprocessed and the ones that contained potential ligation junctions were truncated to keep the longest piece without a junction sequence, including the restriction site sequence. The ligation junction is the concatenation of two filled-in restriction sites: AAGCTAGCTT for HindIII, which cuts at A|AGCTT; CCATGCATGG for NcoI, which cuts at C|CATGG; GATCGATC for MboI and DpnII that cut at GATC|. The pre-truncation step was performed using a Python function, where the only inputs required are the fastq files and the restriction enzyme (HindIII, NcoI, MboI or DpnII). If a custom restriction enzyme is used, the function allows to input the correspondent ligation junction sequence. A log file is automatically generated with the information about the percentage of reads that have been truncated, and also the length distribution of the truncated reads is plotted in a histogram. Besides pre-truncation, we did not perform any read quality filtering before mapping. After this, mapping was performed using Bowtie 2 with custom parameters, and read pairs were mapped independently on the reference genome to avoid any proximity constraint. After mapping, each fastq file has a corresponding SAM file and log file with statistics of alignment. Then, unmapped reads were discarded and only high quality mapped reads (MAPQ ≥ 30) were kept in each SAM file (Yaffe and Tanay, 2011) using SAMtools. After filtering, only paired reads from the SAM files were kept (Unix code) and finally they were converted to BAM format. The two BAM files derived from alignment and processing were not merged because HiFive requires separated BAM files, one per each read of the read pairs.

### Fragment end (FEND) file

The fragment-end (FEND) file is computed by scanning the reference genome for the restriction sites (RS) using Bowtie 2 with the −k flag set to 8000000 (*hg38*: HindIII, 1.72 RS; MboI, 5.02 million RS - *mm10*: HindIII, 1.69 million RS; MboI, 2.99 million RS). The −k argument changes Bowtie 2 research behavior. By default, Bowtie 2 searches for distinct, valid alignments for each read. When it finds a valid alignment, it continues looking for alignments that are nearly as good or better and the best alignment found is reported. When −k <int> is specified, Bowtie 2 searches for at most <int> distinct, valid alignments for each read. The search terminates when it cannot find more distinct valid alignments, or when it finds <int>, which happens first. In this case, −k 8000000 assured to cover all the cutting sites throughout the genome, for every restriction enzyme. The alignment produced a SAM file, which was converted to BAM (SAMtools) and then BED format (BEDTtools). In order to correct biases according to Yaffe and Tanay, 2011, information about GC content and sequence uniqueness of the fragments was needed. Given the high time complexity of this job, the pipeline has been optimized for parallelized computation using multiple threads. The first step was to split the FEND file into separated BED files, one per each chromosome. Then, several steps were performed to add first the GC content information, and second the mappability score. The information about the GC percentage was taken from the UCSC table browser (track: GC percent), and saved to separate txt files, one per each chromosome. Specifically, the GC content of the 200 bp upstream and downstream of each restriction site was calculated, and this was done using 24 parallel threads for *hg38*, one per chromosome. The processor we used was the Xeon E5-2697 v3 (2.60 GHz) and for the FEND file of MboI on hg38 (5.02 million restriction sites), the computation time was decreased from 100 hours (single thread) to 9 hours. To compute the fragment end mappability score the entire genome was split in into 50 bp artificial reads starting every 10 bp (Python) to produce a fastq file. The artificial reads were then mapped back to the genome using Bowtie 2. For each fragment end, the mappability score was then calculated as the portion of artificial reads mapped with MAPQ > 30 within a 500 bp window upstream and downstream of each restriction site. Same as for the GC content, this job was done using 24 parallel threads and this reduced the computation time the same (9 hours against 100 hours with a single thread). After this, fragment ends with a mappability score less than 0.5 (either upstream and downstream) were discarded.

### Data normalization and visualization

Data normalization was performed using the Python package HiFive (Sauria et al., 2015). Both technical biases (spurious ligation products, fragment length, GC content, mappability score) and biological features (TSSs and CTCF-bound sites) were considered in our pipeline. To remove spurious ligation products, we filtered out paired reads whose total distance from the nearest restriction sites was greater than 500 bp. In addition, PCR duplicates were removed and reads with ends mapping to the same fragment and reads with ends mapping to adjacent fragments on opposite strands were also excluded, to consider the possibility of incomplete restriction enzyme digestion and fragment circularization. To take into account of TSSs and CTCF-bound sites and the way they influence the contact frequency upstream and downstream of them, fragment ends within a distance of 500 kb are excluded in the learning correction parameter model. This allows to normalize data without confounding technical biases with features associated to biological-relevant structures. Fragment length, GC content, mappability score and inter-fragment distance biases are handled and removed using the Binning algorithm, which uses the same probabilistic model as Yaffe and Tanay, 2011. Fragment length, GC content, mappability score and inter-fragment distance ranges are divided into 20 bins, such that each bin contains the same number of fragments. For the optimization process of the correction matrices by likelihood maximization, we used a learning threshold of 1 and a maximum number of iterations of 1000.

Heatmaps and histograms were generated and plotted using the Python libraries Matplotlib and Matplotlib.pyplot. The plotting function for normalized contacts contains a parameter which allows to customize the colormap, either by choosing from one of those listed here https://matplotlib.org/examples/color/colormaps_reference.html or even by generating a custom one by inserting the colors to be used into a list. In addition, there is the possibility to select a cut-off as a percentile of the contact counts. Values above this cutoff will be plotted in a different color respect to the colormap, which can be chosen by the user as well.

### Directionality Index and TADs computation

Given a division of the genome into 40 kb bins, we quantified the observed Directionality Index (DI) using the following formula from Dixon et al., 2012:

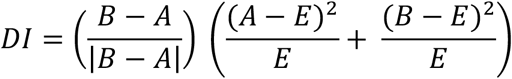

where A is the number of reads that map from a given 40 kb bin to the upstream 2 Mb region, B is the number of reads that map from a given 40 kb bin to the downstream 2 Mb region and E is the expected number of contacts for each bin and it equals to 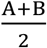. Therefore, the FEND normalized contact data at a bin size of 40 kb is needed to compute the DI. The detection region of 2 Mb for upstream or downstream biases corresponds to 50 bins (2 Mb / 40 kb = 50 bins).

We used a Hidden Markov Model (HMM) based on the Directionality Index to identify biased states. To perform the HMM we used the Python package hmmlearn. Specifically, we built a model with three biased states corresponding to downstream bias, upstream bias and no bias. The sequence of emissions corresponds to the observed DI values and transition matrix, emission matrix and initial state sequence are unknown. We have three types of emissions named as 1, 2, 0 in the model and corresponding to a positive (1), negative (2) or zero (0) value of the observed DI. in our analysis, we associated to the emission ‘0’ all the absolute DI values under a threshold of 0.4. We initialized transition and emission matrices with the same values of 0.3 for the probabilities to transit to a different state or emission respectively (values outside the diagonal), and 0.4 for the probabilities of remaining in the same state or observing the same emission (values in the diagonal). So, first we estimated the model parameters and then the most probable sequence of states using the Viterbi algorithm. Biased states were then exploited to calculate topological domain coordinates. According to Dixon et al., 2012, a domain is initiated at the beginning of a single downstream biased state. The domain is continuous throughout any consecutive downstream biased states and ends when the last in a series of upstream biased states is reached.

### Contact map storage

HiCtool allows to generate and save contact matrices at the resolution defined by the user. In our pipeline, we processed Hi-C data with a bin size of 40 kb to allow topological domains analysis (Dixon et al., 2012). Already at this resolution, contact matrices contain several million of elements per each chromosome, requiring big storage space and relatively high data saving and loading time. To address this problem, we proposed a way to parse the data based on the fact that contact maps are symmetric (contacts between loci *i* and *j* are the same than those between loci *j* and *i*) and usually sparse, since most of the elements are zeros, and this property is stronger with the decreasing of the bin size. Given these two properties, it is not needed to save mirrored data and moreover it would be useful to “compress” the zero data within the matrices. To accomplish this, first we selected only the upper-triangular part of the contact matrices (including the diagonal) and we reshaped the data by rows to form a vector. After that, we replaced all the consecutive zeros in the vector with a “0” followed by the number of zeros that are repeated consecutively; all the non-zero elements are left as they are. Finally, the data are saved in a txt file (**Figure S1**). As mentioned before, at higher resolutions (low bin sizes) contact matrices show more zeros, meaning that the advantage given by this data format would be more remarkable in terms of storage usage and computation time.

**Figure S1.**
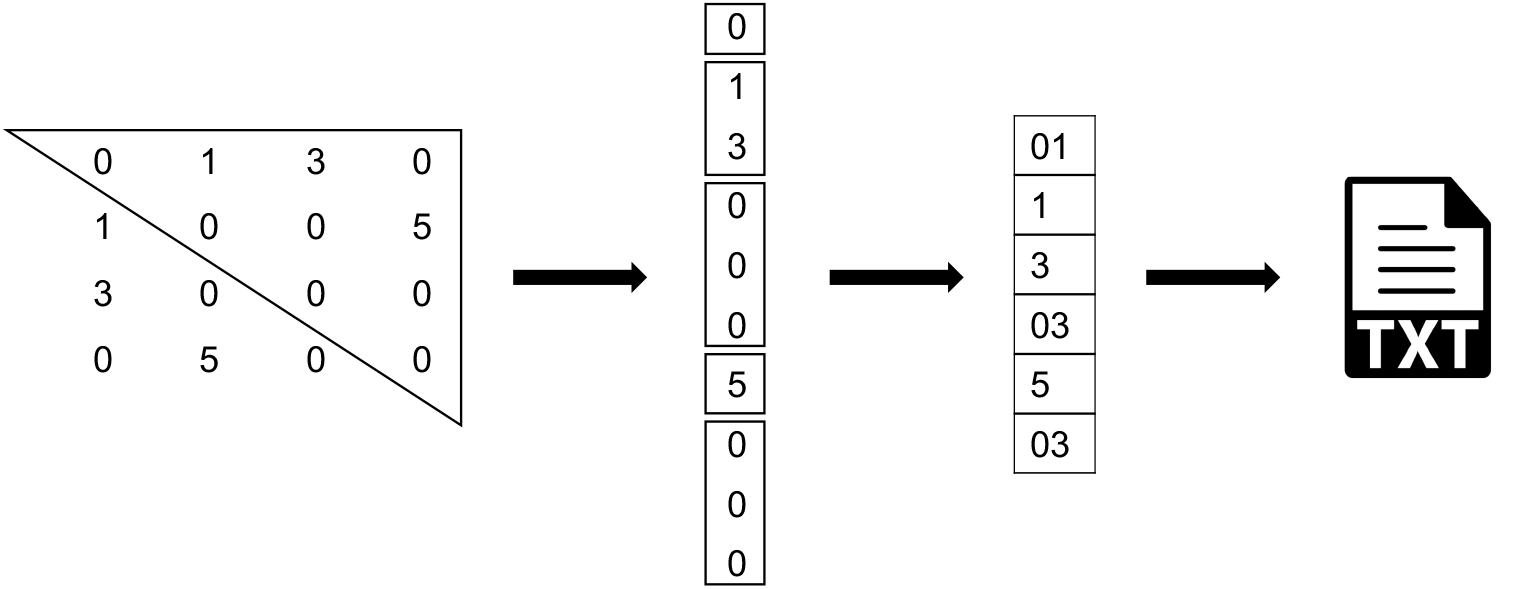
Contact map storage workflow. This is a simplified example where an intra-chromosomal contact matrix is represented by a 4×4 symmetric and sparse matrix. 1) The upper-triangular part of the matrix is selected (including the diagonal). 2) Data are reshaped to form a vector. 3) All the consecutive zeros are replaced with a “0” followed by the number of zeros that are repeated consecutively. 4) Data are saved into a txt file.

### Software requirements

This is the list of software that are required to use GITAR: Python (>2.7), Bowtie 2, BEDTools, SAMTools, SRA Toolkit. The Python libraries needed are: Numpy, Scipy, Math, Matplotlib, Matplotlib.pyplot, Csv, Pybedtools, Pandas, Multiprocessing, Biopython. Additional Python packages are: HiFive and Hmmlearn. HiFive is used to normalize contact data, while Hmmlearn serves for the Hidden Markov Model to calculate the biased states used to extract topological domain coordinates.

